# Loss of clathrin heavy chain enhances actin-dependent stiffness of mouse embryonic stem cells

**DOI:** 10.1101/2020.05.10.086579

**Authors:** Ridim D Mote, Jyoti Yadav, Surya Bansi Singh, Mahak Tiwari, Shivprasad Patil, Deepa Subramanyam

**Author notes:** These authors contributed equally to this study (Ridim D Mote, Jyoti Yadav).

## Abstract

Mouse embryonic stem cells (mESCs) display unique mechanical properties, including low cell stiffness, and specific responses to features of the underlying substratum. Using atomic force microscopy (AFM), we demonstrate that mESCs lacking the clathrin heavy chain (*Cltc*), display higher Young’s modulus, indicative of greater cellular stiffness, in comparison to WT mESCs. We have previously shown that mESCs lacking *Cltc* display a loss of pluripotency, and an initiation of differentiation. The increased stiffness observed in these cells was accompanied by the presence of actin stress fibres and accumulation of the inactive, phosphorylated, actin binding protein, Cofilin. Treatment of *Cltc* knockdown mESCs with actin polymerization inhibitors resulted in a decrease in the Young’s modulus, to values similar to those obtained with WT mESCs. However, the expression profile of pluripotency factors was not rescued. This indicates that a restoration of mechanical properties, through modulation of the actin cytoskeleton, may not always be accompanied by a change in the expression of critical transcription factors that regulate the state of a stem cell, and that this may be dependent on the presence of active endocytosis in a cell.

## Introduction

Early mammalian development is a complex process where different molecular mechanisms and signaling pathways regulate the choice of cell fate. Embryonic stem cells (ESCs) derived from the 3.5 dpc blastocyst serve as a fantastic model system to study early cell fate decisions, as they have the ability to differentiate into all cell lineages [1]. This property of ESCs is governed by numerous factors including transcriptional networks involving molecules such as Oct4, Sox2, Nanog etc. [2], [3], chromatin modifiers such as DNA methyltransferases and histone methyltransferases [4]–[7], endocytic pathways [8], [9], and even mechanical properties [10]–[12]. The core pluripotency transcription factors Oct4, Sox2 and Nanog regulate early embryonic development, and are essential to maintain the identity of ESCs [13]–[15], with loss of Oct4 [13] and Sox2 [14] resulting in early embryonic lethality.

In addition to the role of transcription factors and epigenetic regulators, a growing body of work implicates the process of intracellular trafficking and endocytosis in regulating the fate of embryonic stem cells. Specific pathways, such as the clathrin-mediated endocytic pathway, are essential for maintaining the pluripotent state of mouse ESCs [8]. Other pathways, such as those involving Caveolin, are largely absent in mESCs [9]. Endocytic proteins such as Asrij, are also important for maintaining the pluripotent state of ESCs [16]. Furthermore, it has been demonstrated that the expression of endocytic genes is altered during human somatic cell reprogramming [17]. Endocytosis is also affected by mechanical properties of the cell, such as membrane stiffness, with increasing stiffness resulting in an inhibition of vesicular trafficking [18]–[20]. The effect of stiffness on endocytosis can be countered by an active involvement of actin at endocytic sites [21], [22].

Mechanical properties have also been shown to regulate the pluripotency of ESCs, with cell stiffness or elasticity being one of the major mechanical parameters governing cell fate. Atomic force microscopic (AFM) analysis has been carried out on pluripotent mESCs and early differentiating mESCs, where early differentiating mESCs showed a two to three-fold increase in their elastic modulus compared to naive mESCs [11]. Cell stiffness is governed by a number of cellular properties, of which the actin cytoskeleton plays a major role [23]. Study of the actin cytoskeleton in mESCs has revealed it to be a low-density meshwork, possessing larger pore size, and independent of myosin in mESCs compared to differentiated cells [24]. This study also revealed that the mechanical properties of mESCs may not be completely dictated by the state of the actin cortex. Inhibition of actin polymerization in mESCs has also been shown to result in a decrease in differentiation towards the mesodermal lineage coupled with an increase in differentiation towards the endodermal lineage [25]. Together, these studies reveal that the differentiation of ESCs is a tightly regulated process and requires an intricate interplay between mechanical, cytoskeletal and transcriptional factors.

Previous work from our lab has demonstrated an important role for the Clathrin heavy chain (*Cltc*) in maintaining the pluripotent state of mESCs. CLTC is an integral part of the clathrin coat in clathrin-mediated endocytosis (CME). Knockdown of *Cltc* in mESCs resulted in loss of CME and pluripotency of mESCs, resulting in an initiation of differentiation [8]. Previous studies have shown that differentiating mESCs have a high stiffness and higher Young’s modulus as compared to pluripotent mESCs [11]. As a follow-up to our previous study [8], we asked whether the loss of CME was also accompanied by changes in the stiffness of these cells, similar to other differentiating cells. We also attempted to understand the underlying molecular mechanism that may be involved.

Using atomic force microscopy (AFM), we measured for the first time the Young’s modulus of live mESCs lacking *Cltc* plated on matrigel using a spherical bead attached to a cantilever. We showed that cells lacking *Cltc* have a higher Young’s modulus compared to wild-type (WT) mESCs. We further demonstrated that mESCs lacking *Cltc* displayed an enhancement in the presence of actin stress fibres in cells, which were largely absent in WT mESCs. This was also accompanied by an elevated expression of the inactive, phosphorylated form of the actin depolymerizing protein, Cofilin, resulting in the presence of stable actin filaments. Treatment of *Cltc* knockdown mESCs with the actin polymerization inhibitors, Latrunculin A and Cytochalasin D, resulted in a rescue of cellular stiffness, with cells reverting to a state closer to WT mESCs with respect to mechanical properties. However, the expression profile of pluripotency factors was not rescued, and continued to resemble that of a differentiating cell, indicating that alterations in the actin cytosksleton may be able to regulate pluripotency only in the presence of active CME in mESCs. Together these results suggest that the pluripotent state is an amalgamation of both mechanical and molecular properties, that function together to influence the state of a cell. Additionally, a change in a single readout may not be sufficient to completely predict or alter the state of a cell.

## Results

### *Cltc* knockdown results in increased cell stiffness

Clathrin-mediated endocytosis (CME) is a type of vesicular transport in which receptors are internalized into intracellular structures called endosomes with the help of the coat protein, clathrin [26]. We have previously shown that knockdown of the clathrin heavy chain (*Cltc*) in mESCs, resulted in a decreased expression of pluripotency markers and an increased expression of differentiation markers of all three germ layers [8]. However, the mechanical properties of mESCs under these conditions remained to be interrogated. *Cltc* was knocked down in mESCs using lentiviral-based shRNA-mediated knockdown (Supp. Fig. 1A and 1B and [8]. Two of the three shRNAs (shCltc1 and shCltc3) showed significant knockdown of *Cltc* in mESCs compared to cells infected with the lentivirus expressing the scrambled shRNA, resulting in a significant decrease in protein levels of CLTC (Supp. Fig. 1A and 1B), and were used for all further experiments.

A novel method employing a tipless cantilever with stiffness of 0.03-0.09N/m with a spherical glass bead of diameter 5µm attached to the end was used for all AFM measurements (Fig. 1A, Supp. Fig. 2) (see Materials and methods). mESCs (either expressing a scrambled shRNA, or *Cltc* shRNA) were plated on Matrigel for all AFM measurements. Matrigel supports the growth and maintenance of a variety of cells [27], and has been previously used as a substrate for the AFM analysis of ESCs [10]. In AFM investigations, the spatial resolution depends on the sharpness of the tip. Sharp pyramidal tips are generally used for measurements of mechanical response at a subcellular level on components such as the cytoplasm and nucleus, combined with high resolution imaging. Spherical beads attached to tipless cantilevers are generally used for deformations and resulting stress modulation of inhomogeneous surfaces and are used to investigate the response of the cell as a whole [28]. The spherical tips provide more contact area for measurement, thereby resulting in the reduction of strain on the membrane preventing cell damage. AFMs measure the Young’s modulus of a wide range of biological materials from subcellular features to tissues and organisms and also provide unprecedented spatial resolution, and correctly measure the relative changes in stiffness before and after certain molecules are knocked down.

**Figure 1:**
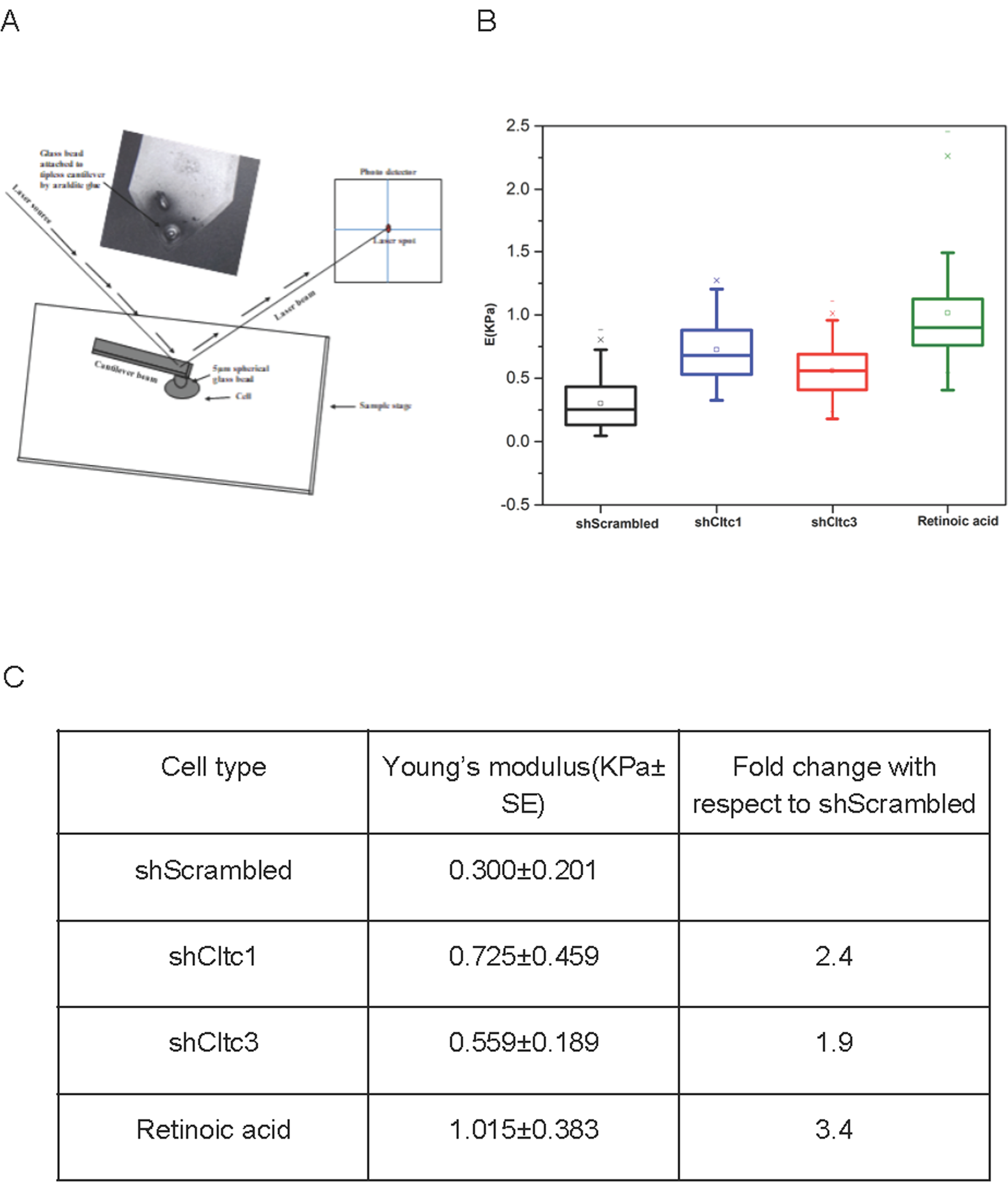
*Cltc* knockdown results in increased cell stiffness: A) Schematic of AFM cantilever interacting with cell. The inset shows an image of the cantilever attached with a 5 µm bead. B) Young’s modulus (E) of shScrambled, shCltc1, shCltc3 and retinoic acid treated mESCs. C) The table shows the apparent Young’s modulus (E) of undifferentiated (shScrambled), *Cltc* knockdown (shCltc1, shCltc3), and retinoic acid treated mESCs.

The inherent approximation in all these studies is that the glass-tip contact is non-deformable and hence has infinite stiffness. As a result all the reported values of stiffness using AFM may have a systematic error in their absolute values. However, this assumption is largely valid for measurements on cells and tissues for which the Young’s modulus is of the order of few KPa. The supplementary materials show representative force curves on cells with respect to glass and matrigel (Supp. Fig. 2).

To measure the Young’s modulus of matrigel, a 10×10*µ*m area was selected and measurements were taken at 5 different points. Hertz mechanics was used to determine the Young’s modulus [29]. The Young’s modulus of matrigel was found to be in the range of 2-3KPa. These values are higher than those reported earlier [10], [30], and can be attibuted to the differences in protocols adopted for preparing the gel and also the size of the microspheres used in the AFM measurement.

AFM was used to determine the Young’s modulus of live mESCs infected with lentiviruses expressing either shScrambled, shCltc1 or shCltc3 plated on matrigel-coated coverslips. Measurements were made on 10 cells for each sample. Young’s modulus (E) for mESCs expressing scrambled shRNA was 0.300±0.201KPa whereas E for cells expressing shCltc1 was 0.725±0.459KPa, and for shCltc3 was 0.559±0.189KPa, indicating higher stiffness in *Cltc* knockdown mESCs (Fig. 1B, 1C, Supp Fig. 2). Measurements were also made from mESCs treated with retinoic acid for 48 hrs to induce differentiation, resulting in Young’s modulus of 1.015±0.383KPa, indicating much greater stiffness (Fig. 1B, 1C, Supp Fig. 2), with the raw data clearly revealing a variation in relative stiffness of cells under different knock-down conditions (Supp. Fig. 2). Our results demonstrate that the Young’s modulus increased by 2.4 fold in shCltc1, 1.9 fold in shCltc3, and by 3.4 fold in mESCs grown in retinoic acid with respect to wild-type mESCs, indicating an increase in cell stiffness upon loss of CME and/or subsequent differentiation.

### Actin cytoskeleton reorganization upon *Cltc* knockdown in mESCs

The actin cytoskeleton is the one of the major regulators of cellular stiffness as it provides mechanical stability to adherent cells [24], [31]. Phalloidin staining revealed that actin stress fibres were predominantly present in *Cltc* knockdown mESCs compared to scrambled cells (Fig. 2A). Actin filaments are dynamic structures and are involved in multiple cellular processes such as cell migration, cell division, endocytosis etc [32]. Actin filament dynamics are regulated by a number of actin-binding proteins. Among the actin-binding proteins, Actin-depolymerizing factors (ADF) or destrin and cofilin family proteins are involved in the depolymerization of actin filaments. LIM-kinases (LIMK1 and LIMK2) and related testicular protein kinases (TESK1 andTESK2 in mammals) are known to inhibit the activity of cofilin. These kinases phosphorylate cofilin at serine 3 which inhibits cofilin’s actin depolymerizing activity [32]. Hence, we investigated the status of cofilin phosphorylation in *Cltc* knockdown mESCs. Phosphorylation of cofilin at serine 3 residue was higher in *Cltc* knockdown mESCs as compared to scrambled shRNA-treated mESCs (Fig. 2B,C). This indicates that cofilin is inactivated by serine 3 phosphorylation in *Cltc* knockdown mESCs, consistent with the observation of abundant actin stress fibres upon knockdown of *Cltc* in mESCs (Fig. 2A). Treatment of scrambled as well as *Cltc* knockdown mESCs with an inhibitor of actin polymerization, Latrunculin A in *Cltc* knockdown mESCs resulted in a discernible loss of actin fibres (Fig. 2D). Together, these results indicate that the loss of *Cltc* results in an increase in the presence of actin stress fibres and that this can be reversed by the action of actin polymerization inhibitors.

**Figure 2:**
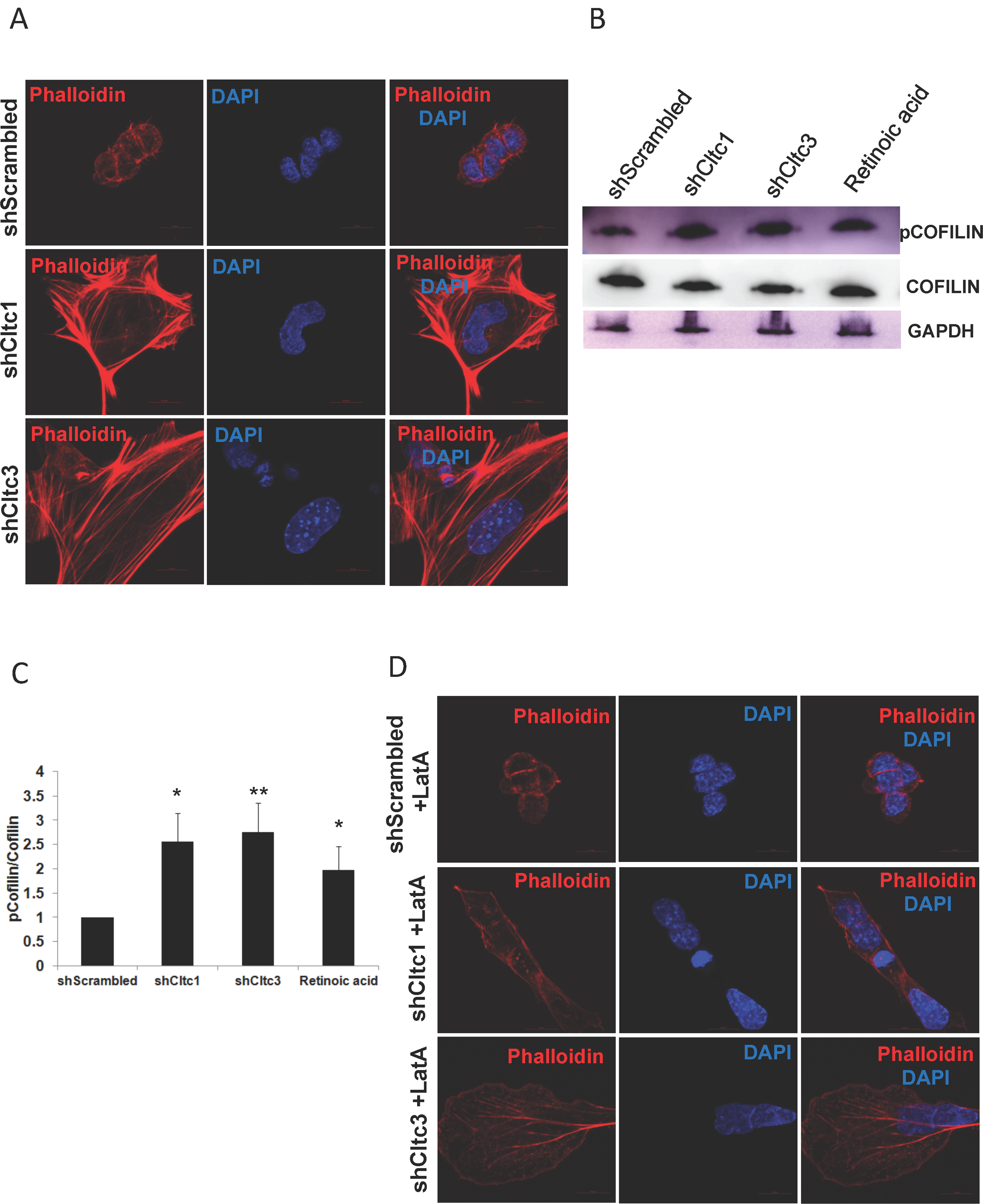
Changes in actin cytoskeleton architecture upon *Cltc* knockdown in mESCs: A) Representative confocal micrographs showing actin filaments stained with Phalloidin in shScrambled, shCltc1, shCltc3 and retinoic acid treated V6.5 mESCs. The scale bar represents 10µm. N=15. B) Western blot showing expression of pCOFILIN, COFILIN and GAPDH in shScrambled, shCltc1, shCltc3 V6.5 mESCs (N=3); and C) Quantitation of pCofilin/Cofilin in shScrambled, shCltc1, shCltc3 V6.5 mESCs (N=3). For all experiments, error bars represent mean ± S.D for experiments in triplicates (N = 3). *p < 0.05; **p < 0.01 by Students T-test. D) Representative confocal micrographs showing actin filaments in shScrambled, shCltc1, shCltc3 mESCs treated with Latrunculin A (0.1 uM) for 12hrs and stained using Phalloidin. The scale bar represents 10µm. N=15.

### F-actin depolymerizing agents reduces the stiffness of *Cltc* knockdown mESCs without rescuing the expression of pluripotency markers

To further validate whether the actin cytoskeleton was indeed the major regulator of cell stiffness, and for the differentiation observed in *Cltc* deficient mESCs, we treated scrambled as well as *Cltc* knockdown mESCs with inhibitors of actin polymerization, Latrunculin A and Cytochalasin D. Upon treatment with Latrunculin A, the Young’s modulus for shScrambled mESCs was 0.208±0.07KPa (Fig. 3A, 3C), indicating a decrease in stiffness compared to untreated shScrambled mESCs (Fig. 1B), similar to what has been previously reported [24]. A reduction in the Young’s modulus was also observed for shCltc1 (0.221±0.094KPa) and for shCltc3 (0.232±0.115KPa) upon Latrunculin A treatment (Fig. 3A, 3C), consistent with our observation of a decrease in stress fibres (Fig. 2D). Similarly upon treatment with Cytochalasin D, the Young’s modulus for shCltc1 and shCltc3 reduced to 0.186±0.100KPa and 0.292±0.087KPa, respectively, and was comparable to the Young’s modulus for treated shScrambled mESCs (0.260±0.102KPa) (Fig. 3B, 3C). Since treatment of *Cltc* knockdown mESCs with Latrunculin A or Cytochalasin D resulted in a decrease in stress fibres and cell stiffness to levels similar to those of pluripotent WT mESCs, we next asked whether the expression of pluripotency markers was also restored to levels seen in WT mESCs. *Cltc* knockdown in mESCs showed decreased expression of pluripotency markers (Fig. 3D), as previously reported [8]. However, treatment with Latrunculin A or Cytochalasin D did not result in a significant change in the expression of pluripotency markers (Fig. 3D), suggesting that this may not be possible in the absence of active CME. Together these results indicate that while the actin cytoskeleton is largely involved in the regulation of cellular stiffness, its modulation may not result in an alteration of the pluripotency network under conditions where CME is not functional.

**Figure 3:**
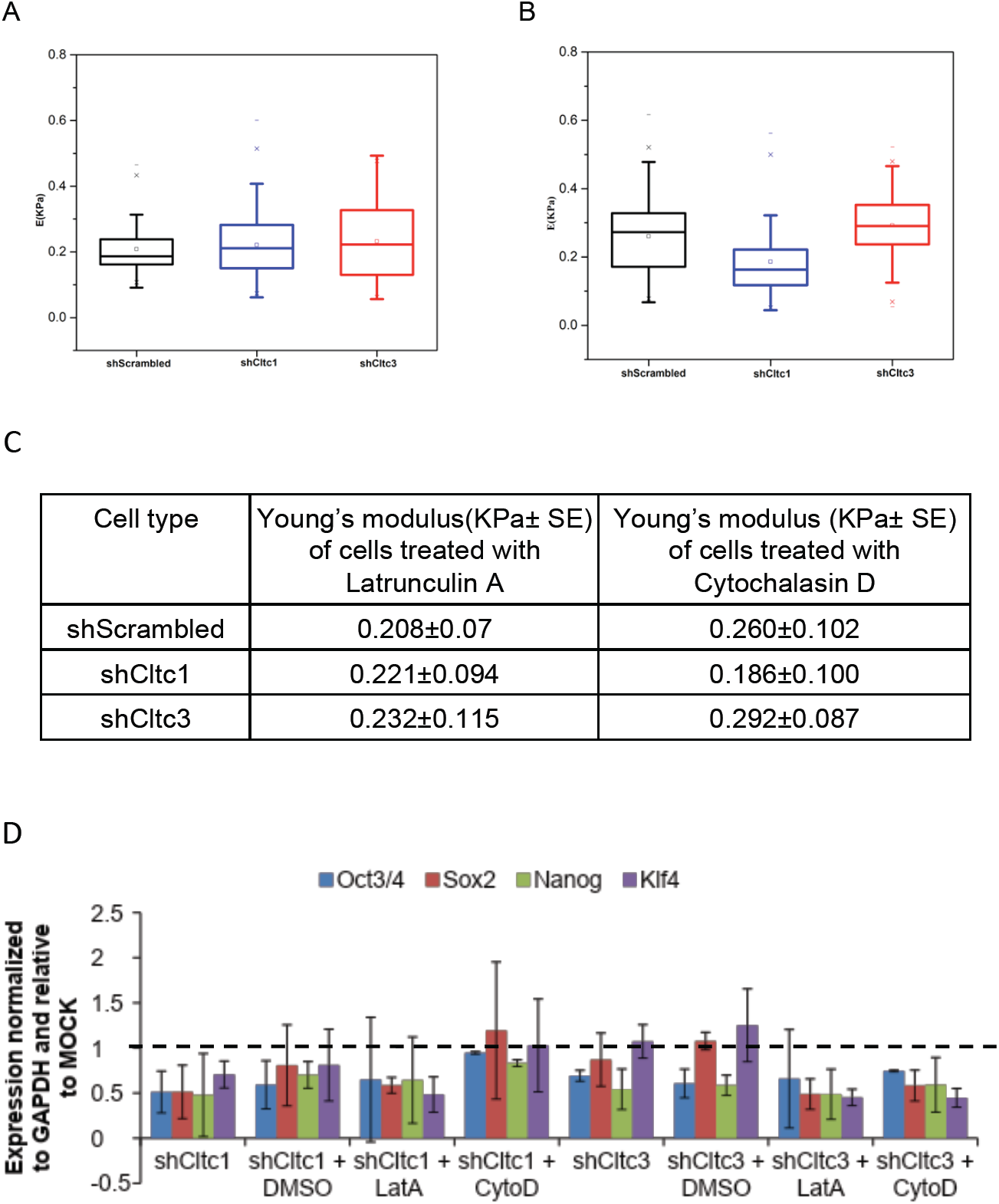
Actin depolymerizing agents reduce the stiffness of *Cltc* knockdown mESCs: A) and B) Young’s modulus (E) of shScrambled, shCltc1, shCltc3 treated V6.5 mESCs treated with Latrunculin A (A), and Cytochalasin D (B). C) Table showing Young’s modulus (E) of shScrambled, shCltc1, shCltc3 treated mESCs. D) RT-qPCR analysis of pluripotency markers in shScrambled, shCltc1 and shCltc3. Bar graph showing the expression of pluripotency markers in mESCs under indicated conditions relative to the relevant shScrambled control. Control is shown as a dotted line at 1. Error bars represent range for experiments in duplicate (N = 2).

## Discussion

We have recently demonstrated that the loss of CME results in decreased expression of pluripotency factors in mESCs, accompanied by an increase in the expression of differentiation markers [8]. Here, we show that mESCs lacking CME display greater stiffness or Young’s modulus (E), as demonstrated by AFM measurements (Fig. 1). Previous studies have also reported that the stiffness of early differentiating ESCs is higher compared to undifferentiated ESCs as determined by AFM [11]. Our data also demonstrates that while the organization of the actin cytoskeleton in mESCs lacking *Cltc*, is largely responsible for their higher mechanical stiffness compared to their WT counterparts, it may not be solely responsible for modulating the level of expression of pluripotency markers (Fig. 2, 3).

ESCs are a potential source of cells for regenerative medicine which have applications for therapeutic purposes in a variety of diseases. Treatment for these diseases require efficient protocols for terminal differentiation and ultimately purification of these cells. Classically, researchers have concentrated on a handful of markers to decide whether a cell is pluripotent or has undergone differentiation to give rise to a specialized cell type. More recently, it has become increasingly obvious that other factors contribute by varying degrees to the pluripotent state of a stem cell. These include endocytic pathways and proteins, mechanical properties, in addition to transcription factors and epigenetic modifiers. Recent reports have described using mechanical phenotyping as an label-free and efficient technique for large-scale purification of cells [33]. However, a better understanding of the interplay between mechanical properties and expression of stem cell-specific transcription factors is required before mechanical phenotyping can be used for large-scale purification of cells.

Cellular stiffness has been shown to be accompanied by an increased polymerization of the actin cytoskeleton. Studies using super-resolution microscopy have demonstrated that mESCs have a low-density meshwork of F-actin, with large pore sizes compared to differentiated cells [24]. Our data shows that the loss of CME in mESCs resulted in the presence of stress fibres, which were largely absent in WT mESCs. These cells also displayed an enhanced level of the inactive, phosphorylated form of the actin-binding protein cofilin, in *Cltc* knockdown and RA treated ESCs. This phosphorylation results in the inhibition of cofilin activity, resulting in a block in actin depolymerization, causing an abundance of actin stress fibres upon *Cltc* knockdown in ESCs as compared to WT ESCs. Previous reports have demonstrated that knocking down the actin capping protein Capzb also resulted in increased stiffness [24]. The increase in stiffness of mESCs upon knockdown of *Cltc* could be rescued by treatment with inhibitors of actin polymerization, Latrunculin A or Cytochalasin D. Previous reports have also shown that the treatment of mESCs with Cytochalasin D, resulted in a decrease in cellular stiffness [24]. Xia et al also demonstrated that while myosin activity did not play a predominant role in regulating the elastic modulus of ESCs, an interplay of Formin and Arp2/3 activity were involved [24]. However, the effect of these mechanical changes on the expression of pluripotency factors was not examined. We observed that even though treatment of *Cltc* knockdown mESCs with actin polymerization inhibitors reduced their stiffness to levels similar to WT mESCs, the expression of pluripotency markers could not be rescued under these conditions, suggesting that active intracellular transport through the clathrin pathway may be a critical requirement in attaining pluripotency. An initiation of CME may be essential for the transport of molecules that ultimately regulate the pluripotency network of a stem cell. Our results also suggest that a rescue of mechanical properties need not necessarily always reflect a change in the transcriptional network of an mESC.

Furthermore this may also suggest that an inherent hierarchy may exist with respect to specific events that dictate when a cell achieves pluripotency. A pluripotent ESC is thus the result of a complex interplay between many different molecular, intracellular and mechanical players.

## Materials and methods

### Mouse embryonic stem cell culture

V6.5 mouse embryonic stem cells (mESCs) were cultured on tissue culture plates (Corning) coated with 0.2% gelatin. mESCs were maintained in Knockout DMEM supplemented with 15% fetal bovine serum, 0.1 mM beta mercaptoethanol, 2mM L-glutamine, 0.1 mM nonessential amino acids, 5000 U/ml penicillin/streptomycin, and 1000 U/ml LIF (ESC medium). Cells were passaged using trypsin, every 3 days.

### Lentivirus packaging and infection

Lentiviral vectors pLKO.1-shScrambled, pLKO.1-shCltc1/ shCltc2/ shCltc3 were individually cotransfected with psPAX2 (ADDGENE NO. 12260), pMD2.G (ADDGENE NO. 12259) in HEK293T cells using FuGENE HD (Promega E2311). pLKO.1-shCltc1, shCltc2 and shCltc3 were obtained from the shRNA Resource Centre, Indian Institute of Science, Bangalore. Viral supernatants were harvested and used for infection 60 hours post transfection. 1.5 × 10^5^ mESCs were infected with packaged lentiviruses with polybrene in suspension, followed by plating on gelatin-coated plates (Corning CLS3516). The following day, lentivirus-containing media was removed and ESC media was added to the cells. 24 hours later, cells were trypsinized and 3 × 10^4^ cells were plated on matrigel (Corning 356234) coated 22mm coverslips in ESC media containing puromycin (1ug/ml). AFM experiments were performed on live cells 12 hours post plating.

### Inhibitor treatment

5 × 10^4^ mESCs of shScrambled, shCltc1 and 3 were plated in 0.2% gelatin coated 24 well plate. After 12hrs of plating, cells were treated with the following inhibitors of actin polymerization: Latrunculin A (Sigma L5163) at 0.1 uM final concentration for 12 hrs; and Cytochalasin D (Sigma C8273) at 0.2 uM final concentration for 12 hrs. After 12 hrs, samples were processed for RNA isolation. Retinoic acid (RA) was added to mECSs at a concentration of 10^−7^M in the absence of LIF.

### RNA isolation and Real-time PCR

Total RNA was isolated from mESCs using TRIzol and quantified using a Nanodrop spectrophotometer. 1ug of total RNA was used to synthesize complementary DNA (cDNA) using Verso cDNA synthesis Kit from Thermo Scientific (#AB-1453/A). Gene-specific primers for RT-qPCR were designed using IDT software (sequences provided in Supp. Table 1). ABI Power SYBR Green PCR master mix was used for quantitative RT–PCR reactions. ABI qPCR system, 7900 HT was used to perform quantitative RT–PCR reactions.

### Western blotting

Protein lysates were prepared from cells using RIPA buffer containing proteinase inhibitors and PMSF. Protein concentration was determined using Bradford reagent. Equal concentration of total protein lysates were subjected to SDS-PAGE electrophoresis followed by transfer to PVDF membrane. After transfer, the membrane was blocked in 5% BSA. Post blocking, the membrane was incubated with the appropriate primary antibody at 4°C overnight with gentle rocking. Next day the blot was washed twice for 10 min using Tris-buffered saline (1X TBS) containing 0.1% Tween-20 (TBS-T). Membranes were incubated with an HRP-conjugated secondary antibody (1:1000) for 1 hour at room temperature. Thermo Scientific SuperSignal West Femto Maximum Sensitivity Substrate (Cat. no. 34095) reagent was added to the membranes, and images were captured post-exposure using a chemi-doc system (GE Healthcare, AI600). Western blot images were quantified using ImageJ software. Statistical analysis: For all experiments, error bars represent mean ± S.D for experiments in triplicates (N = 3). *p < 0.05; **p < 0.01; ***p < 0.001 by Students T-test.

### Immunocytochemistry and imaging

2 × 10^4^ mESCs were cultured on glass coverslips coated with matrigel in a 24 well plate. Following day, cells were washed with PBS, fixed with 4% paraformaldehyde at room temperature for 20 min, and permeabilized with 0.1% Triton X-100 in PBS for 5 min. After blocking in 5% bovine serum albumin in PBS for 1 h, cells were incubated with Phalloidin Alexa Fluor 568 (1:400) for 30 min. After 1 wash with 1X PBS, nuclei were stained with DAPI. After 2 washes, coverslips were mounted onto glass slides using VECTASHIELD (Vector Laboratories). Images were acquired using a Nikon Ti Eclipse confocal microscope with Plan Apo 60x oil objective (NA 1.4) at 512 × 512 pixels and 8 bit resolution.

### Bead attachment on cantilever

A tipless cantilever with stiffness of 0.03-0.09N/m was used for AFM measurement. A spherical glass bead with diameter 5µm was attached to the end of the cantilever. The attachment of the bead on the cantilever was done using micromanipulation available with AFM. An approximate equal amount of araldite adhesive (locally sourced) and hardener glue was taken and mixed properly. With the help of a sharp toothpick, small dots were made on a glass slide. Using the servo control of AFM, the cantilever was brought down on the araldite glue. Since the bead was of very small size, a very small amount of glue was picked up on the cantilever, and the cantilever was again lowered down on the glass slide in order to remove the excess glue. After that the cantilever was lowered down onto the bead. Cantilever was maintained under positive load and after 5 minutes, the cantilever was pulled back along with the bead. Before measurements, the cantilever was calibrated each time on the glass coverslip.

### Cell probing using AFM

A thin layer of matrigel was coated on 22mm glass coverslips. Stiffness of matrigel was in the range of 2-3KPa. All measurements were done on live cells only. Indentation studies were performed on 10 control (undifferentiated) mESCs and 10 treated cells (*Cltc* knockdown or RA-treated). The cantilever was aligned in the middle of the cell and the indentation force curve profile was then recorded. The approach velocity was 2µm/sec and force curve resolution was 2048 data points per second.

### AFM analysis

Assuming the glass-glass contact to be infinitely stiff compared to the glass-cell contact - a reasonable assumption since it is 10,000 times stiffer, the slope of the curve in the contact region for glass and matrigel coated glass is nearly one implying no deformation (Supp. Fig. 2). The slope of the curve on cells is much less suggesting a certain amount of deformation. We used glass-glass contact for calibration of deflection sensitivity, and the subtraction of cantilever deflection from the push given by the piezo extension yields deformation in the tissue. The force is calculated by multiplying the cantilever deflections by its stiffness. The force versus deformation curve is then fitted with Hertz model.

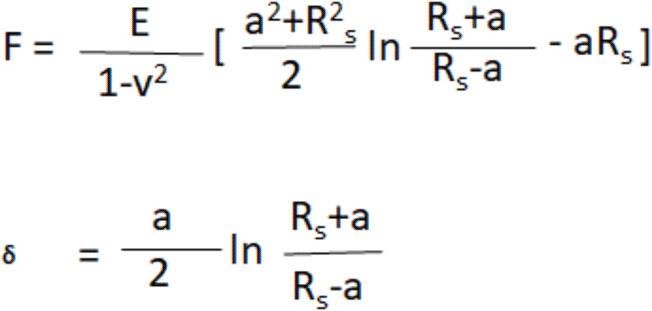

Where F is measured by the cantilever possessing the bead, which is pressed against the cell. R is the bead radius, delta is the deformation in the cell, E is Young’s modulus and v is the Poisson ratio. Hertz mechanics describes the mechanics of cells that are probed with spherical tip and works for non-adhesive elastic contacts.

## Supplementary Information

**Supplementary Figure 1:**
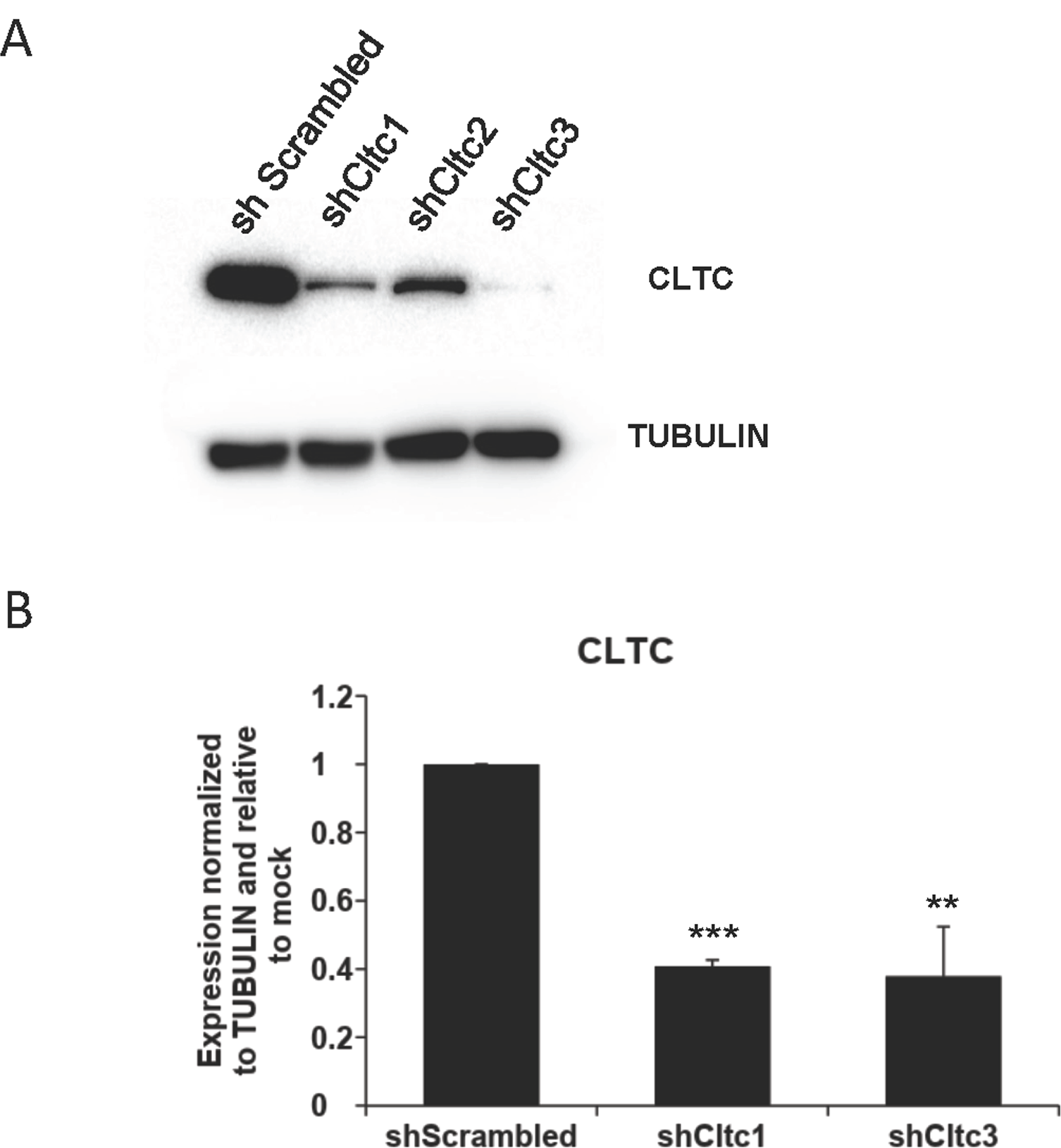
*Cltc* knockdown in mESCs: A) Western blot showing expression of CLTC and TUBULIN in shScrambled, shCltc1, shCltc3 V6.5 mESCs (N=3), and; B) Quantitation of CLTC knockdown in shScrambled, shCltc1, shCltc3 V6.5 mESCs (N=3). For all experiments, error bars represent mean ± S.D for experiments in triplicates (N = 3). *p < 0.05; **p < 0.01; ***p < 0.001 by Students T-test.

**Supplementary Figure 2:**
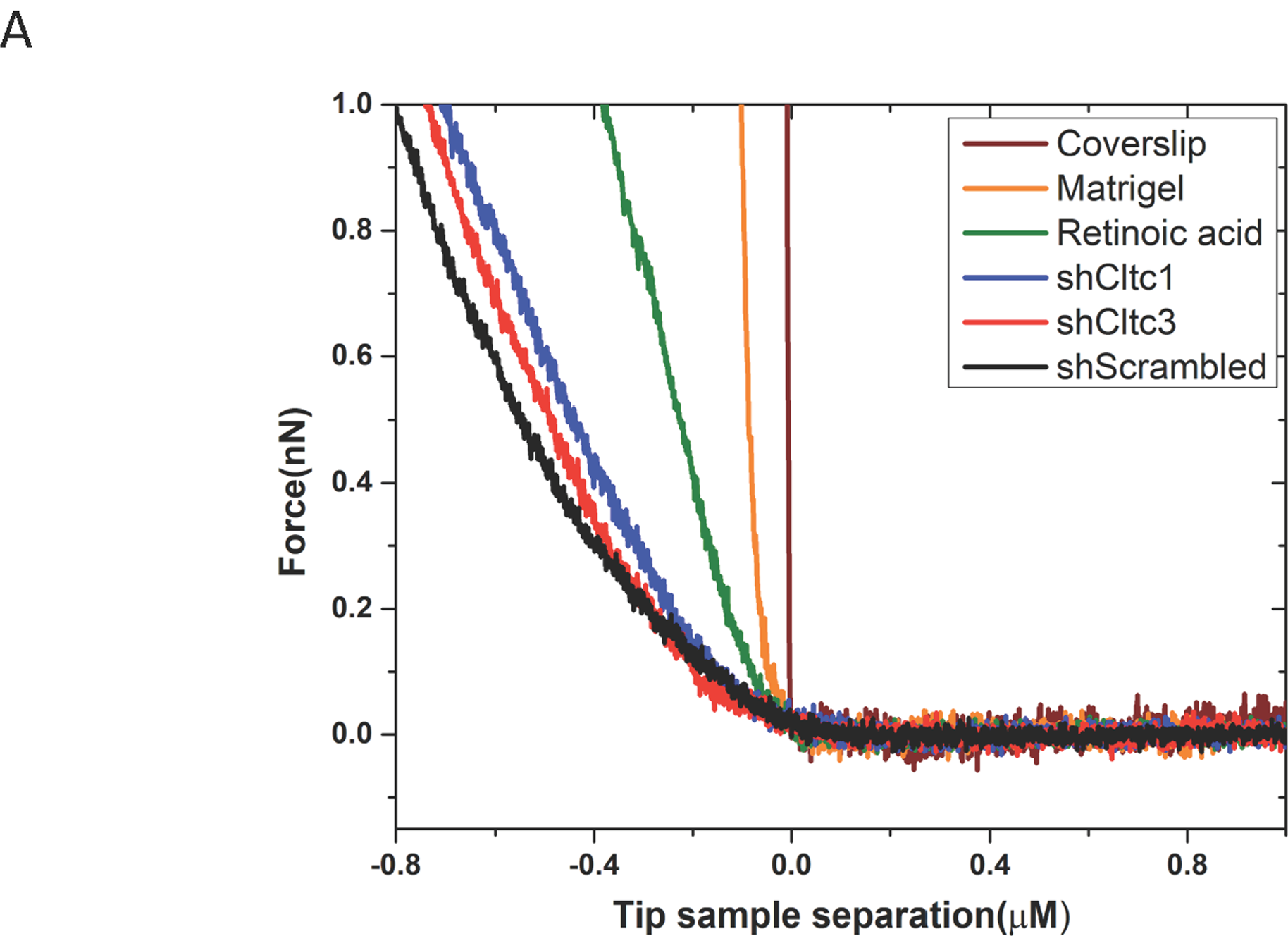
Representative force curves on samples: Representative force curves on all samples plotted together. This data is used to estimate the Young’s modulus reported in Fig. 1 and Fig. 3 in the main text. The raw data clearly reveals variation in stiffness in mESCs after various treatments and knockdowns. The glass cover-slip is a reference and the tip-glass contact is assumed to be non-deforming.

**Supplementary Table 1:**
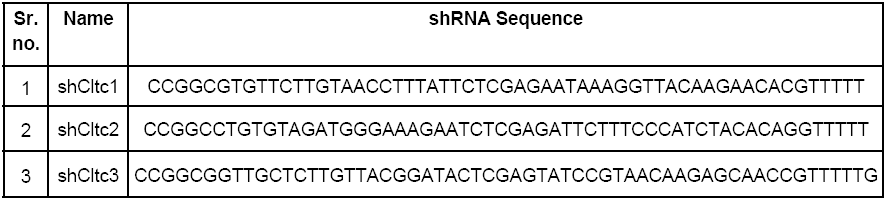
A) Table showing shRNA sequence for different *Cltc* clones. B) List of primers used in this study.

## Acknowledgements

This work was supported by funds to D.S. from the Wellcome Trust – Department of Biotechnology (DBT) India Alliance (Intermediate Fellowship-IA/I/12/1/500507), and the Department of Biotechnology (BT/PR30450/MED/31/399/2018). S.P acknowledges the Wellcome Trust-Department of Biotechnology (DBT) India Alliance (Intermediate Fellowship 500172/Z/09/Z). J.Y. is a recipient of a Senior Research Fellowship from University Grants Commission, India; S.B.S is a recipient of a Senior Research Fellowship from the Department of Biotechnology, India; M.T. is a recipient of a Junior Research Fellowship from University Grants Commission, India. We thank members of the Subramanyam lab for constructive discussion.

## Author contributions

D.S and R.D.M. conceived and designed the study. R.D.M. contributed to experiments described in Fig. 1B-C 2A-D, 3A-D, Supp. Figs. 1A-B. J.Y and S.P. did the acquisition and analysis for experiments described in Fig. 1A-C, 3A-C, Supp. Fig. 2. S.B.S helped with imaging for experiments described in Fig. 2A, 2D. M.T contributed to the experiment described in Fig. 3D. D.S, S.P, R.D.M and J.Y wrote the manuscript. All authors reviewed the manuscript.

---

**Supplementary Table 2.**
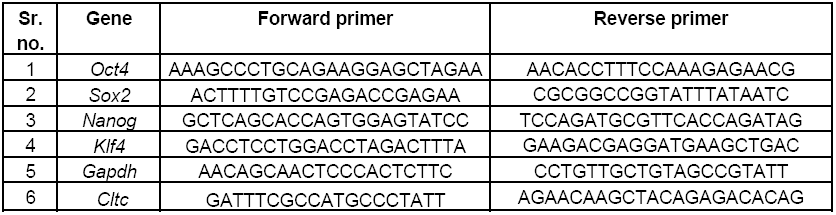

